# SARS-CoV-2 Omicron XBB.1.5 May Be a Variant That Spreads More Widely and Faster Than Other Variants

**DOI:** 10.1101/2023.01.18.524660

**Authors:** Aki Sugano, Haruyuki Kataguchi, Mika Ohta, Yoshiaki Someya, Shigemi Kimura, Yoshimasa Maniwa, Toshihide Tabata, Yutaka Takaoka

## Abstract

In this research, we aimed to predict the relative risk of the recent new variants of severe acute respiratory syndrome coronavirus 2 (SARS-CoV-2) on the basis of our previous research. We first performed molecular docking simulation analyses of the spike proteins with human angiotensin-converting enzyme 2 (ACE2) to determine the binding affinities to human cells of three new variants of SARS-CoV-2: Omicron BQ.1, XBB, and XBB.1.5 We then investigated the three variants to discover the evolutionary distance of the spike protein gene (S gene) from the Wuhan, Omicron BA.1, and Omicron BA.4/5 variants, to understand the changes in the S gene.

The results indicated that the XBB.1.5 variant had the highest binding affinity of the spike protein with ACE2 and the longest evolutionary distance of the S gene. This *in silico* evidence suggested that the XBB.1.5 variant may produce infections that spread more widely and faster than can infections of preexisting variants.

## 1. Introduction

Infection by the new Omicron variant of severe acute respiratory syndrome coronavirus 2 (SARS-CoV-2) has been an ongoing epidemic disease consisting of different successive variants. In early 2023, Omicron BQ.1, XBB, and XBB.1.5 were discovered in patients and were thought to present a particular risk inasmuch as these variants may suggest a coming epidemic. Previously, we reported the *in silico* infectivity of SARS-CoV-2 variants—Alpha, Beta, Gamma, Delta, and Omicron BA.1, BA.2, and BA.2.75—as the ratio per Wuhan variant and the absolute evolutionary distance of the spike protein gene (S gene) between Wuhan and each variant [1]. In our research here, we report the predicted risks for Omicron BQ.1, XBB, and XBB.1.5, which were recently recognized as being causes of epidemic diseases. For this purpose, we utilized the analytic methods of the docking simulation and the evolutionary distance that we had developed in our previous research [1-3].

## 2. Materials and methods

### 2.1 Determination of the absolute evolutionary distances of the variant S genes, and docking simulation for the affinities of the different spike proteins with angiotensin-converting enzyme 2 (ACE2)

We analyzed the absolute evolutionary distances of the S gene from the Wuhan, Omicron BA.1, and Omicron BA.4/5 variants for the following variants —Alpha, Beta, Gamma, Delta, and Omicron BA.1, BA.2, BA.4/5, BA.2.75, BQ.1, XBB, and XBB.1.5—via the ClustalW program [4] and the FastTree program [5]. We obtained the sequences of the S gene by searching the NCBI database (MN908947 for Wuhan, OW519813 for Alpha, and OM791325 for BA.1) or the EpiCoV database of GISAID (https://gisaid.org) for the complete sequences of the S gene (EPI_ISL_5142896 for Beta, EPI_ISL_14534452 for Gamma, EPI_ISL_4572746 for Delta, EPI_ISL_13580480 for BA.2, EPI_ISL_13304903 for BA.4/5, and EPI_ISL_14572678 for BA.2.75, EPI_ISL_15638667 for BQ.1, EPI_ISL_16344389 for XBB, EPI_ISL_15802393 for XBB.1.5).

We obtained the amino acid substitution information (Table 1) of the spike proteins mostly from the CoVariants website (https://covariants.org). We then used the amino acid sequences for the analyses of the three-dimensional structures of each variant spike protein according to our previous research [1].

**Table 1:**
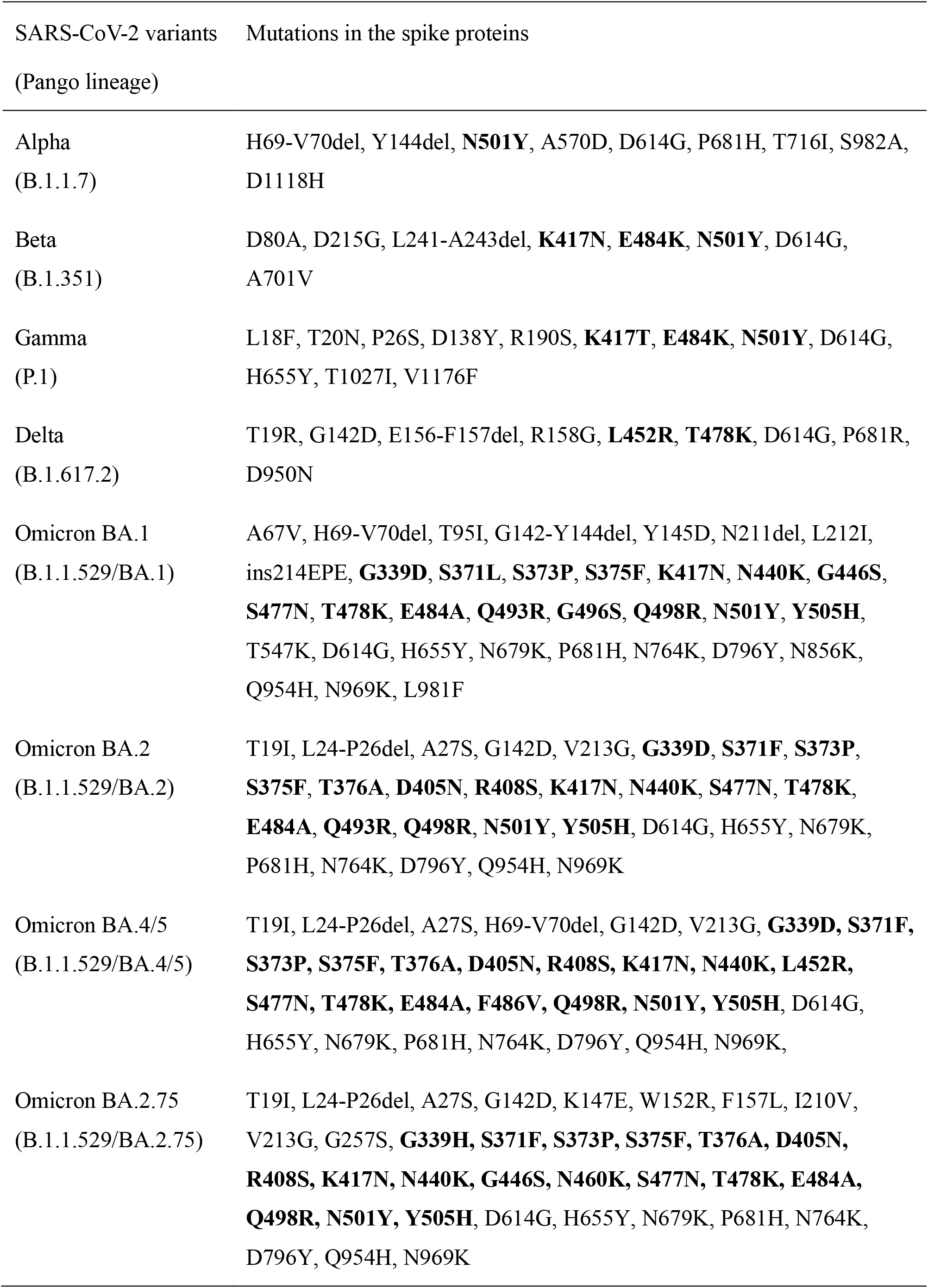
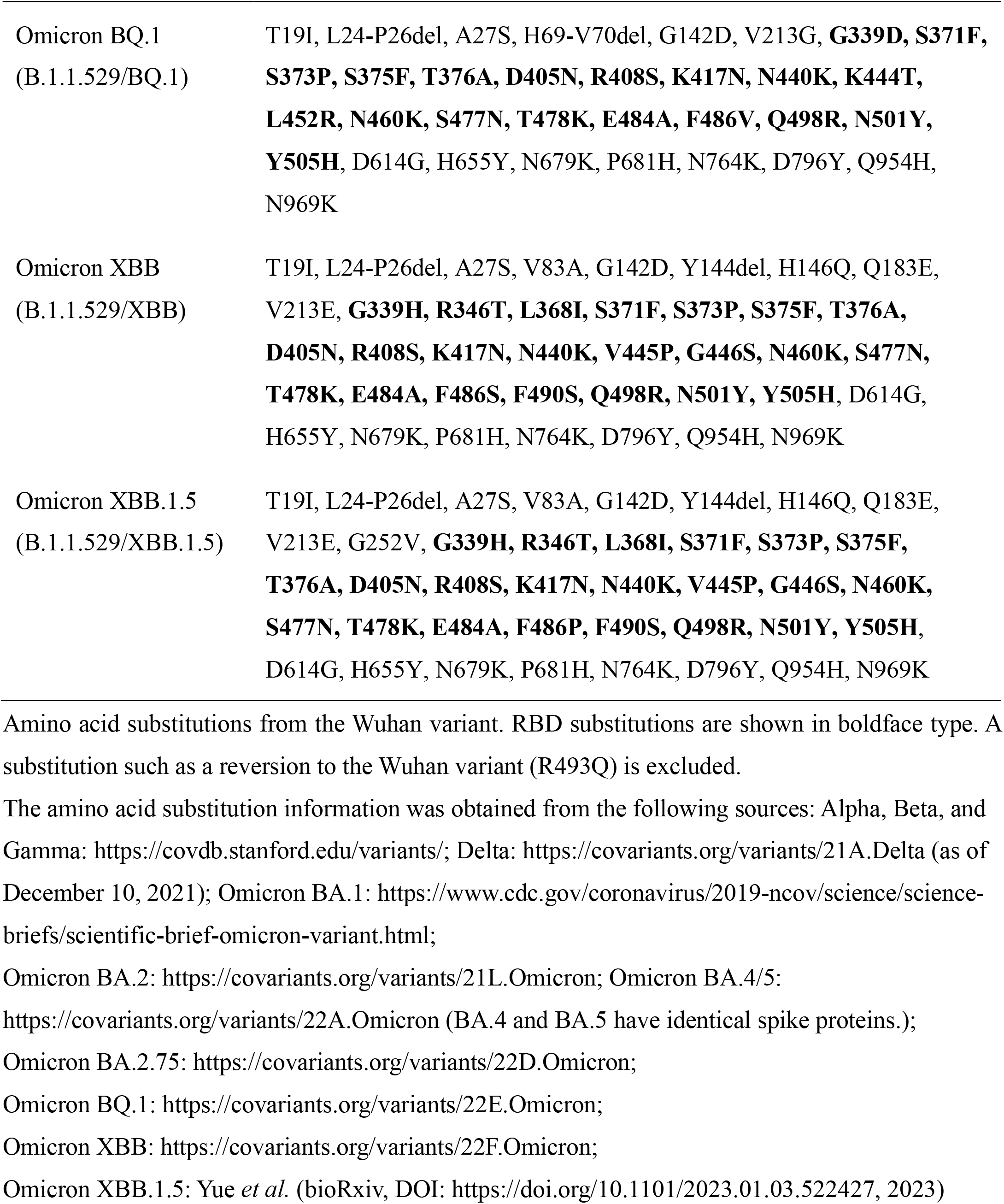
Amino acid substitutions in spike proteins of the different SARS-CoV-2 variants.

| SARS-CoV-2 variants<br>(Pango lineage) | Mutations in the spike proteins |
| --- | --- |
| Alpha<br>(B.1.1.7) | H69-V70del, Y144del, **N501Y**, A570D, D614G, P681H, T716I, S982A, D1118H |
| Beta<br>(B.1.351) | D80A, D215G, L241-A243del, **K417N**, **E484K**, **N501Y**, D614G, A701V |
| Gamma<br>(P.1) | L18F, T20N, P26S, D138Y, R190S, **K417T**, **E484K**, **N501Y**, D614G, H655Y, T1027I, V1176F |
| Delta<br>(B.1.617.2) | T19R, G142D, E156-F157del, R158G, **L452R**, **T478K**, D614G, P681R, D950N |
| Omicron BA.1<br>(B.1.1.529/BA.1) | A67V, H69-V70del, T95I, G142-Y144del, Y145D, N211del, L212I, ins214EPE, **G339D**, **S371L**, **S373P**, **S375F**, **K417N**, **N440K**, **G446S**, **S477N**, **T478K**, **E484A**, **Q493R**, **G496S**, **Q498R**, **N501Y**, **Y505H**, T547K, D614G, H655Y, N679K, P681H, N764K, D796Y, N856K, Q954H, N969K, L981F |
| Omicron BA.2<br>(B.1.1.529/BA.2) | T19I, L24-P26del, A27S, G142D, V213G, **G339D**, **S371F**, **S373P**, **S375F**, **T376A**, **D405N**, **R408S**, **K417N**, **N440K**, **S477N**, **T478K**, **E484A**, **Q493R**, **Q498R**, **N501Y**, **Y505H**, D614G, H655Y, N679K, P681H, N764K, D796Y, Q954H, N969K |
| Omicron BA.4/5<br>(B.1.1.529/BA.4/5) | T19I, L24-P26del, A27S, H69-V70del, G142D, V213G, **G339D**, **S371F**, **S373P**, **S375F**, **T376A**, **D405N**, **R408S**, **K417N**, **N440K**, **L452R**, **S477N**, **T478K**, **E484A**, **F486V**, **Q498R**, **N501Y**, **Y505H**, D614G, H655Y, N679K, P681H, N764K, D796Y, Q954H, N969K, |
| Omicron BA.2.75<br>(B.1.1.529/BA.2.75) | T19I, L24-P26del, A27S, G142D, K147E, W152R, F157L, I210V, V213G, G257S, **G339H**, **S371F**, **S373P**, **S375F**, **T376A**, **D405N**, **R408S**, **K417N**, **N440K**, **G446S**, **N460K**, **S477N**, **T478K**, **E484A**, **Q498R**, **N501Y**, **Y505H**, D614G, H655Y, N679K, P681H, N764K, D796Y, Q954H, N969K |
| Omicron BQ.1<br>(B.1.1.529/BQ.1) | T19I, L24-P26del, A27S, H69-V70del, G142D, V213G, **G339D, S371F, S373P, S375F, T376A, D405N, R408S, K417N, N440K, K444T, L452R, N460K, S477N, T478K, E484A, F486V, Q498R, N501Y, Y505H**, D614G, H655Y, N679K, P681H, N764K, D796Y, Q954H, N969K |
| Omicron XBB<br>(B.1.1.529/XBB) | T19I, L24-P26del, A27S, V83A, G142D, Y144del, H146Q, Q183E, V213E, **G339H, R346T, L368I, S371F, S373P, S375F, T376A, D405N, R408S, K417N, N440K, V445P, G446S, N460K, S477N, T478K, E484A, F486S, F490S, Q498R, N501Y, Y505H**, D614G, H655Y, N679K, P681H, N764K, D796Y, Q954H, N969K |
| Omicron XBB.1.5<br>(B.1.1.529/XBB.1.5) | T19I, L24-P26del, A27S, V83A, G142D, Y144del, H146Q, Q183E, V213E, G252V, **G339H, R346T, L368I, S371F, S373P, S375F, T376A, D405N, R408S, K417N, N440K, V445P, G446S, N460K, S477N, T478K, E484A, F486P, F490S, Q498R, N501Y, Y505H**, D614G, H655Y, N679K, P681H, N764K, D796Y, Q954H, N969K |

Amino acid substitutions from the Wuhan variant. RBD substitutions are shown in boldface type. A substitution such as a reversion to the Wuhan variant (R493Q) is excluded.
The amino acid substitution information was obtained from the following sources: Alpha, Beta, and Gamma: https://covdb.stanford.edu/variants/; Delta: https://covariants.org/variants/21A.Delta (as of December 10, 2021); Omicron BA.1: https://www.cdc.gov/coronavirus/2019-ncov/science/science-briefs/scientific-brief-omicron-variant.html;
Omicron BA.2: https://covariants.org/variants/21L.Omicron; Omicron BA.4/5:
https://covariants.org/variants/22A.Omicron (BA.4 and BA.5 have identical spike proteins.);
Omicron BA.2.75: https://covariants.org/variants/22D.Omicron;
Omicron BQ.1: https://covariants.org/variants/22E.Omicron;
Omicron XBB: https://covariants.org/variants/22F.Omicron;
Omicron XBB.1.5: Yue *et al.* (bioRxiv, DOI: https://doi.org/10.1101/2023.01.03.522427, 2023)

To clarify the ability of each variant to enter human cells, we performed additional docking simulation to analyze the binding affinity of the receptor binding domain (RBD) of the spike protein with ACE2 for the three variants—BQ.1, XBB, and XBB.1.5 with the same procedure in our previous research [3]. In this research, we defined the binding affinity as the most stable score in the docking results with the correct binding mode.

## 3. Results

### 3.1. Results of docking of the RBD with the ACE2 protein and absolute evolutionary distances for the S gene variants

Table 2 shows the binding affinities of the RBDs of the spike proteins with human ACE2 as the ratio per Wuhan, which we determined from the docking simulation studies including our previous results [1]. The Omicron XBB.1.5 had almost the highest binding affinity, which leads to a potentially high risk of entering human cells.

**Table 2:** The binding affinities of the spike proteins with ACE2 (ratio per Wuhan variant) and the absolute evolutionary distances of the S gene between the Wuhan variant other variants.

| Variants | Wuhan | Alpha | Beta | Gamma | Delta | Omicron | Omicron | Omicron | Omicron | Omicron | Omicron | Omicron |
| --- | --- | --- | --- | --- | --- | --- | --- | --- | --- | --- | --- | --- |
|  |  |  |  |  |  | BA.1 | BA.2 | BA.4/5 | BA.2.75 | BQ.1 | XBB | XBB.1.5 |
| Pango lineage | B | B.1.1.7 | B.1.351 | P.1 | B.1.617.2 | B.1.1.529/ | B.1.1.529/ | B.1.1.529/ | B.1.1.529/ | B.1.1.529/ | B.1.1.529/ | B.1.1.529/ |
|  |  |  |  |  |  | BA.1 | BA.2 | BA.4/5 | BA.2.75 | BQ.1 | XBB | XBB.1.5 |
| Binding affinity of S protein with ACE2 (ratio per Wuhan) | 1 | 1.18 | 1.23 | 1.31 | 2.10 | 1.55 | 2.46 | 2.15 | 2.90 | 3.09 | 1.89 | 3.04 |
| Absolute evolutionary distance of the S gene (from Wuhan) x 10<sup>-3</sup> | - | 2.06 | 2.06 | 3.54 | 3.24 | 10.68 | 8.29 | 9.17 | 10.95 | 10.06 | 12.44 | 13.03 |
| Absolute evolutionary distance of the S gene (from BA.1) x 10<sup>-3</sup> | - | - | - | - | - | - | 5.60 | 6.48 | 8.24 | 7.36 | 9.71 | 10.29 |
| Absolute evolutionary distance of the S gene (from BA.4/5) x 10<sup>-3</sup> | - | - | - | - | - | - | - | - | 2.97 | 1.02 | 4.44 | 5.02 |

Table 2 also shows the absolute evolutionary distances of the S gene between the Wuhan variant and each of the other variants, as well as the evolutionary distances between those variants and the Omicron BA.1 or BA.4/5 variant. The variants with longer evolutionary distances from Wuhan, Omicron BA.1, and Omicron BA.4/5 suggest that they possess a tendency toward causing more epidemics, as based on our previous research [3].

These data suggest that Omicron XBB.1.5 had a possibly weak vaccine effect because it has a long absolute evolutionary distance from the three variants that constitute the basis for developing the vaccine.

## 4. Discussion

In this research, we chose two factors as indicators of virus infectivity on the basis of our previous report [1] as follows: (1) binding affinities between the RBD of the spike proteins and human ACE2, which indicate the ability of the virus to enter human cells; and (2) the evolutionary distances of the S genes, which reflect the effects of vaccines and the induced neutralizing antibodies. We thus analyzed the binding affinities with ACE2 and the evolutionary distances of the S genes, which were calculated from Wuhan, Omicron BA.1, and Omicron BA.4/5.

The binding affinities of the RBDs in the recent new variant spike proteins with ACE2 were greater than those of the preexisting variants except Omicron XBB. The result suggests that the risk of XBB’s entering human cells is not as high as that of BA.2.75, BQ.1, or XBB.1.5, which can be estimated to be between that of BA.1 and Delta or BA.4/5 variants. The evolutionary distances of the recent new Omicron variants—BQ.1, XBB, and XBB.1.5—indicate the following possibilities: Omicron BQ.1 has a short evolutionary distance from BA.4/5, which suggests that BA.4/5-based vaccines can be effective against this variant; Omicron XBB and XBB.1.5 have long evolutionary distances, which suggests that currently available vaccines have a poor effect against these variants. Omicron XBB.1.5 also showed almost the highest binding affinity of the spike protein with the human ACE2 protein compared with other variants, and the S gene evolutionary distance from the three variants for the current vaccine was the longest. This result suggests that the XBB.1.5 infection can be more widespread than can infections of preexisting variants. Indeed, we previously demonstrated a high correlation between the evolutionary distances or the binding affinities and the reported infectivities [3]. Yue *et al*. reported that, according to Surface Plasmon Resonance analysis, XBB.1.5 demonstrated enhanced receptor-binding affinity [6]. Tamura *et al*. reported that XBB was the most resistant variant to BA.2/5 in infected serum and was better able to enter human cells than was BA.2.75 [7]. In addition, the Centers for Disease Control and Prevention (CDC) provided Nowcast estimates of the proportion of XBB.1.5 that was estimated to account for the largest percentage, 49.1%, of the cases in the US in the week that ended January 21 (updated to 49.5% afterwards), and 61.3% in the week that ended January 28, 2023 (https://covid.cdc.gov/covid-data-tracker/#variant-proportions). These reports were consistent with our *in silico* results.

Although our results do not include data on the risk of exacerbation of SARS-CoV-2, they do indicate the need for great caution in managing XBB.1.5, because the number of severely ill patients will increase along with the number of infected individuals, even if this variant has a low risk of exacerbation.

## 5. Conclusion

We report here that the Omicron XBB.1.5 variant of SARS-CoV-2 has the longest evolutionary distance of the S gene from the Wuhan, Omicron BA.1, and Omicron BA.4/5 and almost the highest binding affinity for spike protein with ACE2 according to docking simulation studies. These results suggest that Omicron XBB.1.5 poses a greater risk in the pandemic than other variants.

## Abbreviations

S gene: spike protein gene
ACE2: angiotensin-converting enzyme 2
SARS-CoV-2: severe acute respiratory syndrome coronavirus 2
RBD: receptor binding domain

## Acknowledgments

We gratefully acknowledge all data contributors, i.e., the authors, their originating laboratories that obtained the specimens, and their submitting laboratories for generating the genetic sequences and metadata for sharing this information via the GISAID Initiative; and all data provided by CoVariants, on which this research is based.

## Author Contributions

Y.T. conceived and designed this research. Y.T., A.S., H.K, and M.O. preformed the analyses and acquired the data. Y.T., A.S., H.K., and M.O. interpreted the data. Y.T. and A.S. wrote the draft, and all authors reviewed and approved the manuscript.

## Ethical Approval Statement

This statement is not applicable because we performed computer analyses by using sequence data obtained from a public database.

## Data Availability

Data that support the findings of this study are available from the corresponding author upon reasonable request, except publicly available data sources.

